# Synthetic Natural Product Inspired Peptides

**DOI:** 10.1101/2021.06.15.448394

**Authors:** Matthew A. Hostetler, Chloe Smith, Samantha Nelson, Zachary Budimir, Ramya Modi, Ian Woolsey, Autumn Frerk, Braden Baker, Jessica Gantt, Elizabeth I. Parkinson

**Author notes:** **Corresponding Author** Elizabeth I. Parkinson, Department of Chemistry and Medicinal Chemistry and Molecular Pharmacology, Purdue University, West Lafayette, IN 47907, USA. Matthew Hostetler, Department of Chemistry, Marshall University, Huntington, West Virginia 25755, United States.

## Abstract

Natural products (NPs) are a bountiful source of bioactive molecules. Unfortunately, discovery of novel bioactive NPs is challenging due to cryptic biosynthetic gene clusters (BGCs), low titers, and arduous purifications. Herein, we describe SNaPP (<u>S</u>ynthetic <u>Na</u>tural <u>P</u>roduct Inspired <u>P</u>eptides), a method for identifying NP-inspired bioactive molecules. SNaPP expedites bioactive molecule discovery by combining bioinformatics predictions of non-ribosomal peptide synthetases (NRPS) with chemical synthesis of the predicted NPs (pNPs). SNaPP utilizes a recently discovered cyclase, the penicillin binding protein (PBP)-like cyclase, as the lynchpin for the development of a library of cyclic peptide pNPs. Analysis of 500 BGCs allowed for identification of 131 novel pNPs. 51 diverse pNPs were synthesized using solid phase peptide synthesis and in-solution cyclization. Antibacterial testing revealed 14 pNPs with antibiotic activity, including activity against multidrug-resistant Gram-negative bacteria. Overall, SNaPP demonstrates the power of combining bioinformatics predictions with chemical synthesis to accelerate the discovery of bioactive molecules.

## INTRODUCTION

Natural products (NPs) have been a bountiful source of medicines including antimicrobials, anticancer agents, antiparasitics, immunosuppressants, as well as many others.^1^ Historically, bacteria have been one of Nature’s most prolific producers of biologically active NPs.^2^ One important class of biologically active bacterial NPs are nonribosomal peptides (NRPs). These peptides are synthesized by modular enzyme complexes known as nonribosomal peptide synthetases (NRPS) and comprise a rich set of structurally diverse NPs, including many clinically used antibiotics such as daptomycin, bacitracin, polymyxin B, and colistin.^3^ Cyclic peptides are an especially important class of NRPs, possessing many favorable pharmacological properties over their linear counterparts.^4–6^ Their relatively large size and structural rigidity allow them to engage challenging targets, including protein-protein interactions.^4,7–9^ Cyclic NRPs are also generally more cell permeable and resistant to proteases compared to linear peptides.^5,10,11^ For this reason, there is great interest in the discovery of additional cyclic NRPs as biological tools and drug leads.

Traditionally, novel NRPs have been discovered by a classical fermentation approach^12^ whereby crude bacterial extracts are screened for biological activity. While this approach has been extremely successful, it is very time consuming. The process of going from a bioactive extract to a completely elucidated structure takes minimally several months and oftentimes over a year. Additionally, each new NP requires optimization of fermentation conditions and purification sequences, thus preventing easy automation of the process. Rediscovery of known NPs is also a major limitation.^13^ Recent advances in whole-genome sequencing and bioinformatics have revealed a vast number of NRPS biosynthetic gene clusters (BGCs) for which no known NP can be attributed.^14^ Harnessing the full biosynthetic potential of these organisms is complicated by the fact that a small fraction (~2%) of bacteria are culturable in the laboratory^2,15^ and many BGCs are transcriptionally inactive (cryptic) under standard laboratory conditions.^14^ Access to the NPs produced via these BGCs often requires heterologous expression or promoter optimization, both of which are very time consuming and frequently unsuccessful.

We hypothesized that we could overcome these difficulties by developing SNaPP (**<u>S</u>**ynthetic **<u>Na</u>**tural **<u>P</u>**roduct Inspired **<u>P</u>**eptides), a method that combines bioinformatics with chemical synthesis. Specifically, the method utilizes 1) bioinformatics tools such as antiSMASH^16^ and PRISM^17^ to predict peptide products formed by NRPS BGCs identified in bacterial genomes and 2) chemical synthesis to access the predicted peptides. This synthesis-first approach has many advantages over traditional fermentation approaches: 1) This approach skips bacterial culture and the need for fermentation optimization, 2) It avoids rediscovery of known NPs by comparison with known BGCs, 3) Products from cryptic BGCs or currently unculturable bacteria can easily be accessed, and 4) Each part of SNaPP from the identification of the BGCs to NP predictions to chemical synthesis is scalable and easily automated, greatly expediting the process.

Others have prepared predicted NRPs by solid-phase peptide synthesis and were successful in the discovery of several biologically active compounds.^18–22^ However, few of these reports has explored the synthesis of predicted cyclic NRPs,^22,23^ despite the fact that nearly 67% of known NRPs possess a cyclic motif.^24,25^ One reason for this observation may be the limited ability of bioinformatics programs to predict how NRPs cyclize. The thioesterase (TE) domain is typically the terminal module of an NRPS and is often responsible for peptide cyclization.^26^ However, TE domains catalyze the production of multiple cyclic motifs including lactams and lactones in head-to-tail or sidechain-to-tail form.^27,28^ Interestingly, numerous NRPS BGCs do not contain a thioesterase domain and instead are released from the NRPS via stand-alone enzymes. Recently, the penicillin binding protein (PBP)-like cyclases have been identified as a novel class of stand-alone NRPS cyclases.^29–31^ PBP-like cyclases have only been found to catalyze cyclization of the C-terminus with the N-terminus to furnish head-to-tail cyclic lactams. Herein we describe our efforts to discover novel antibiotic cyclic peptides via the synthesis of predicted NPs (pNPs) that is guided by the selection of NRPS BGCs neighboring PBP-like cyclases. While these peptides are not intended to be the true NPs, we expect to bias ourselves toward head-to-tail cyclic peptides with interesting bioactivities using this selection criteria.

## RESULTS AND DISCUSSION

### Identification of pNPs

SurE, the PBP-like cyclase that catalyzes the cyclization of the surugamides, is one of the most well studied PBP-like cyclases.^29–32^ *surE* along with the genes encoding the PBP-like cyclases for the NPs ulleungmycin (*ulm16*), desotamide B (*dsaJ*), the mannopeptimycins (*mppK*), the pentaminomycins (*penA*), the noursamycins (*nsm16*), and the curacomycins (KUM80512.1) are all found in close proximity to the NRPS that produces the peptide NP.^33–36^ This co-localization suggests that the genes for these cyclases could be used as a genetic handle for identifying other cyclic NRPs. Our strategy is outlined in **Figure 1**. First, a BlastP^37^ search for SurE was performed and the top 500 hits were analyzed further. The genetic neighborhood for these hits was identified using RODEO.^38^ 396 (79%) of the BGCs had NRPS genes 10 genes or less away. Clusters at the end of a contig or with incomplete records in NCBI (80, ~20%) were removed prior to further analysis. The remaining 316 NRPS containing BGCs were then analyzed using bioinformatics softwares including PRISM 4^17^ and antiSMASH 5.0^16^ to predict the structure of the NRPs. Generally, predictions between the two programs agreed well. Tanimoto analysis of the predictions from PRISM 4 or antiSMASH 5.0 for the 5 known molecules within our dataset compared to their actual structures suggested similar accuracies (**Supplementary Figure 1A**). Additionally, their predictions for uncharacterized BGCs also was similar (**Supplmentary Figure 1B**). We ultimately chose to use the PRISM predictions as the basis for our studies for two major reasons. The first reason is that PRISM is more likely to give a structural prediction, even when the prediction is less sure.^39^ For our purposes, it was more useful to have a prediction, even if it was not necessarily a high confidence prediction. The second reason is that other studies have found that PRISM is better at predicting known NPs compared to antiSMASH when the dataset it larger.^39^ Using PRISM, 140 unique cyclic peptides were identified. Nine of the peptides were previously known NRPs (mannopeptimycin, desotamide B, ulleungmycin, and 6 copies of the surugamide cluster) leaving 131 unique and novel cyclic peptides of varying sizes to explore further (**Figure 2A** and **Supplementary Excel File**).

**Figure 1.**
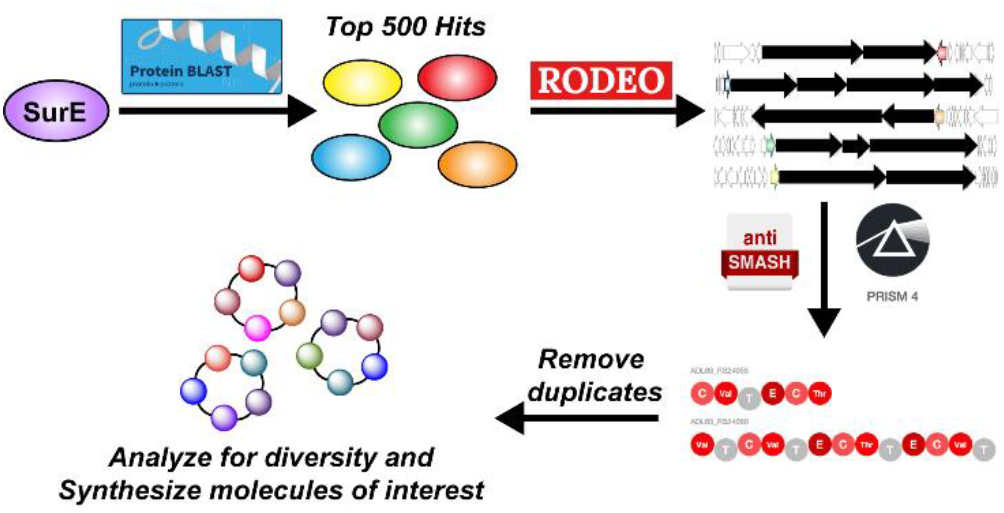
Outline of SNaPP method.

**Figure 2.**
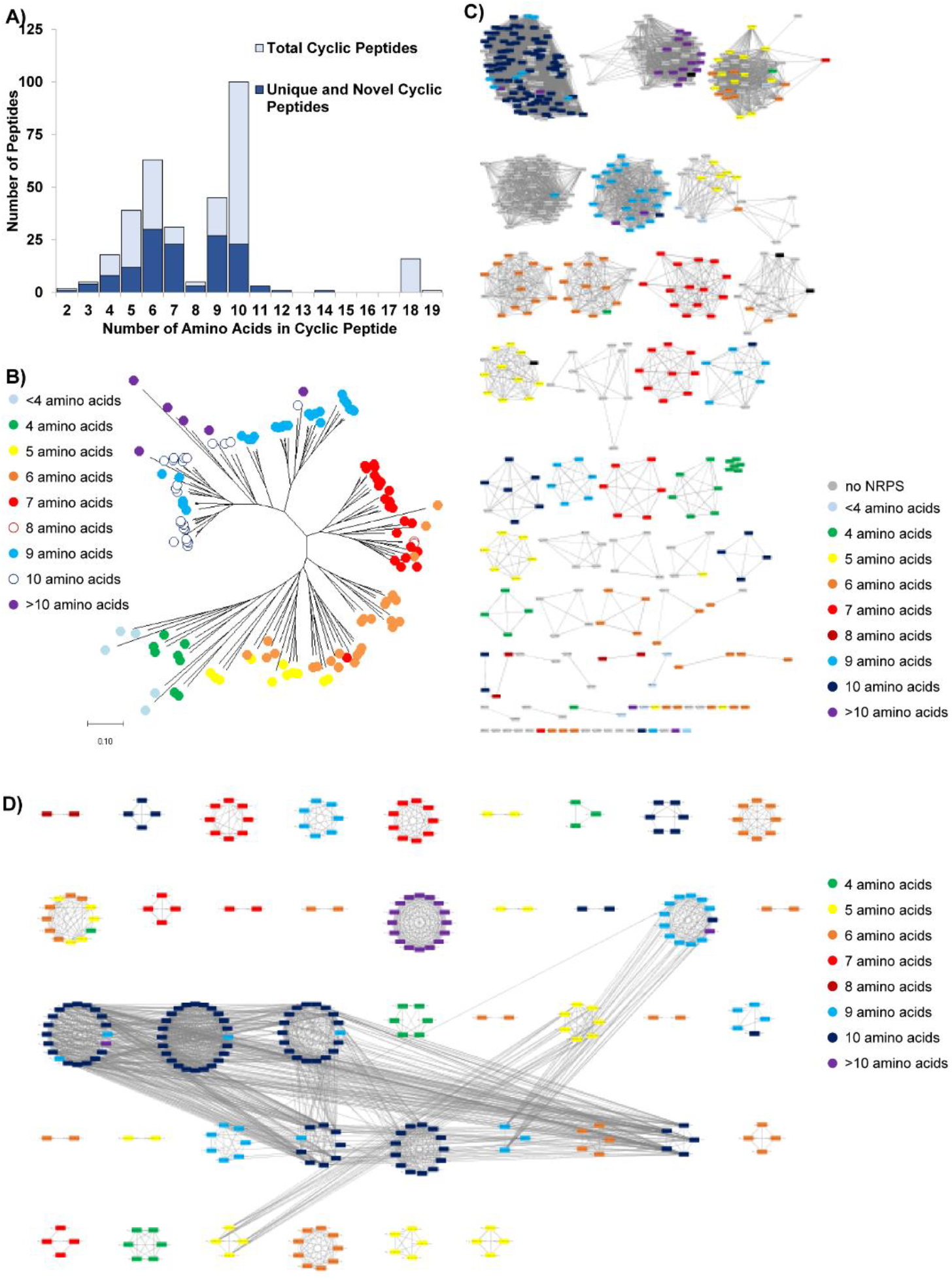
Diversity of pNPs. A) pNPs distribution with total number of cyclic peptides noted in light blue and the number of unique and novel cyclic peptides noted in dark blue. B) Tanimoto similarity data represented in tree form. Details of strains and molecules synthesized can be found in Figure S2. C) Sequence Similarity Network for PBP-like cyclases. The size (number of amino acids) of the predicted cyclic peptide product is indicated by the color of the nodes. D) BiG-SCAPE network of PBP-like cyclase and NRPS containing BGCs. Each circle represents a family (closely related) of BGCs. Branches to other circles indicate clans (more distantly related BGCs). The size (number of amino acids) of the predicted cyclic peptide product is indicated by the color

Previously, Jacques and co-workers found that NRPs vary in size between 2 and 23 amino acids with the most frequent sizes of NRPs being between 7 and 9 amino acids.^40^ While we see many peptides with 7 and 9 amino acids, we see very few with 8 amino acids and instead see a large number of 6 and 10. Additionally, the unnatural amino acid ornithine is predicted much more often than expected. Based on the number of occurrences in the Norine database,^24^ we would expect ~8% of NRPs to contain ornithine. We found that ~70% of our pNPs contain ornithine. It is unclear whether this is due to the prediction software or if ornithine is truly overrepresented in this set of peptides. Interestingly, antiSMASH often predicted glutamine when PRISM predicted ornithine. Another common difference was that antiSMASH would often predict tyrosine when PRISM predicted tryptophan. Given the structural similarity of these amino acids, we are not surprised by these differences.

### Diversity of pNPs

Because the structures of molecules determine their functions, structural diversity is essential for any compound library that will be used for bioactivity screening.^41^ To assess the diversity of the pNPs and determine the best molecules to synthesize for testing, we first used ChemMine Tools^42^ to calculate the Tanimoto coefficients. The Tanimoto coefficients were then used to generate both a heatmap (**Supplementary Excel File**) as well as a tree (**Figure 2B** and **Figure S2A**). Peptides of the same size generally cluster together while still having noticeable structural differences.

Bioinformatics methods were also employed to analyze the diversity of the library. A sequence similarity network (SSN)^43^ of the PBP-like cyclases was generated. The PBP-like cyclases tend to cluster based on the size of their corresponding NRPs, suggesting that PBP-like cyclases might be specific for certain ring sizes (**Figure 2C** and **Figure S3**). Interestingly, occasionally different sizes are predicted within the same ring suggesting that either these cyclases are more flexible or potentially that the NRPS next the cyclase may act in an iterative fashion. We also performed BiGSCAPE analysis^44^ on the BGCs containing the PBP-like cyclases and NRPS genes (**Figure 2D** and **Figure S4**). This analysis revealed 86 NRPS families with an average of 4 BGCs per family. This data, in agreement with the Tanimoto data, confirms a varied set of structures and helps us to design a diverse library.

### Synthesis of a diverse pNP library

51 chemically diverse pNPs were chosen for synthesis (see **Figure S2-4** and **Table S1**). Challenging to access amino acids such as protected enduracididine and hydroxyphenylglycine were replaced with the structurally similar amino acids arginine and phenylglycine, respectively. Linear sequences were prepared using solid-phase peptide synthesis (SPPS) whereby each coupling cycle was promoted using Oxyma and diisopropylcarbodiimide (**Figure S5**).^45,46^ The side chain protected linear peptides were released from the resin using hexafluoroisopropanol. Solution-phase cyclization was achieved using PyBop under high dilution conditions, and global deprotection with trifluoroacetic acid afforded the desired pNPs. In the majority of cases, these products could be precipitated with methyl *tert-*butyl ether to afford cyclic peptide products that were >90-95% pure and thus were sufficiently pure for biological testing. Any products less than 90% pure were further purified by HPLC. Identity of the pNPs was confirmed via mass spectrometry, and purity was determined via analytical HPLC. The entire sequence from pNP prediction through purification can be completed in 7 days and is straightforward enough to be completed by an undergraduate. Additionally, all steps except HPLC purification can easily be accomplished in parallel. Growth of a NP producing organism often takes longer than this, with fermentation optimization, purification, and structure validation regularly exceeding a year.

### Bioactivity testing

Initial compounds were tested for activity against antibiotic sensitive and antibiotic resistant ESKAPE pathogens at concentrations varying between 0.5 and 32 μg/mL using the CLSI microbroth dilution assay.^47^ Any well with greater than 90% death was considered a hit. Overall, 14 hits (MIC ≤ 32 μg/mL) were observed with 4 against Gram-negative organisms (**Figure 3** and **Table S2**), 9 of them being against Gram-positive organisms (**Figure S6A** and **Table S3**) and, and 1 hit against both. This is a very promising hit rate (~30%), especially when compared to other antibiotic discovery programs, which have struggled to find any hits, especially against Gram-negative organisms.^48,49^ An Alamar blue viability assay revealed that these molecules are non-toxic to the A549 non-small cell lung cancer cell line, suggesting they likely have good selectivity for bacterial cells over mammalian cells (**Table S2-3**). Hemolysis assays with human blood revealed that many also had no hemolytic effects at concentrations up to 53 μg/mL (**Table S2-3**) providing strong evidence that they are promising antibiotic leads.

**Figure 3.**
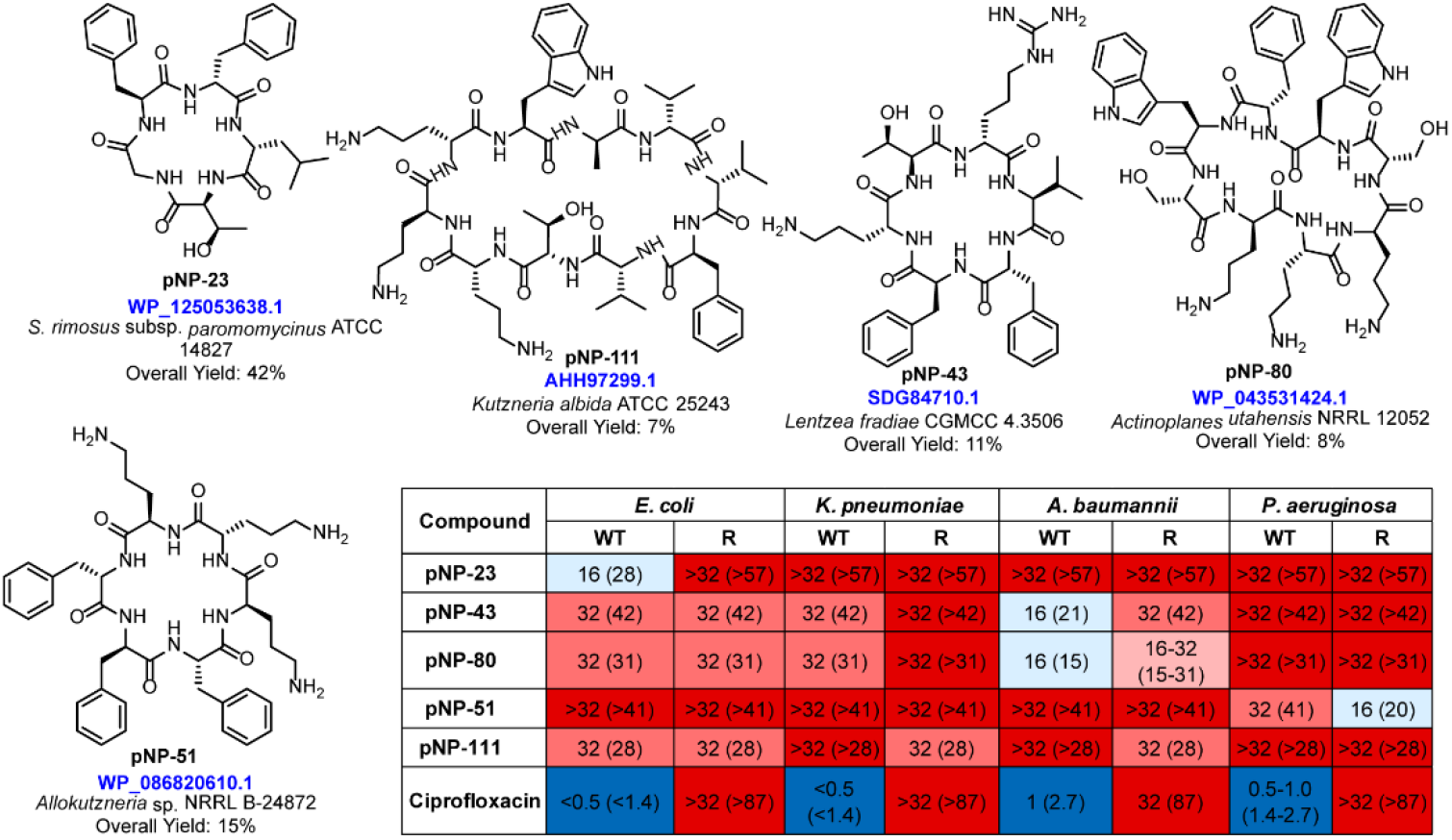
Structures of pNPs that hit against Gram-negative bacteria. Gram-negative bacteria and a table describing their activities. The strains analyzed are described in the materials and methods section. Potencies of hits are given in μg/mL and in parentheses are the potency in μM. WT: wild type; R: antibiotic resistant; Cipro: ciprofloxacin; WT E. coli: ATCC 25922; R E. coli: ATCC BAA-2469; WT K. pneumoniae: ATCC 27736; R K. pneumoniae: ATCC BAA-21469; WT A. baumannii: ATCC 19606; R A. baumannii: KB349; WT P. aeruginosa: PAO1; R P. aeruginosa: PA1000.

### Derivative development and mechanism of action studies

Based on the results described above, we chose to explore derivatives of **pNP-43**, a compound with activity against several Gram-negative bacteria and no observed hemolytic activity or mammalian cell toxicity. **pNP-43** is predicted to be produced by *Lechevalieria fradiae* CGMCC 4.3506, a strain originally isolated from the Wutaishan Mountain in the Shanxi province of China. In addition to the PBP-like cyclase (14) and NRPS genes (15 and 16), the BGC contains genes with high similarity to the enduracididine biosynthetic genes (10-12), providing strong support that enduracididine is incorporated into this cyclic peptide (**Figure 4** and **Table S4**). Structure predictions by PRISM further support this with adenylation domain 6 predicted to load enduracididine. Due to challenges in obtaining enduracididine, we chose to initially substitute enduracididine for the next highest prediction, arginine. When developing derivatives, the arginine was exchanged with amino acids having similar chemical structures including lysine, ornithine, and 2,4-diaminobutyric acid (**pNP-43a-c, Figure S6B**). However, the parent molecule was the most active (**Table S2**). After further examination of the predictions by antiSMASH^16^ and PRISM^17^ (**Table S5**), we chose to develop other derivatives by modifying the amino acid at position 4 (Orn). While ornithine is the number one prediction for amino acid 4, arginine and lysine also scored well thus we chose to incorporate these residues into our derivatives (**pNP-43d-e** in **Figure S6B**). Substituting lysine in place of ornithine at position 4 (**pNP-43d**) resulted in biological activity that was ~4-times more potent against antibiotic resistant *A. baumannii* compared to the initial molecule.

**Figure 4.**
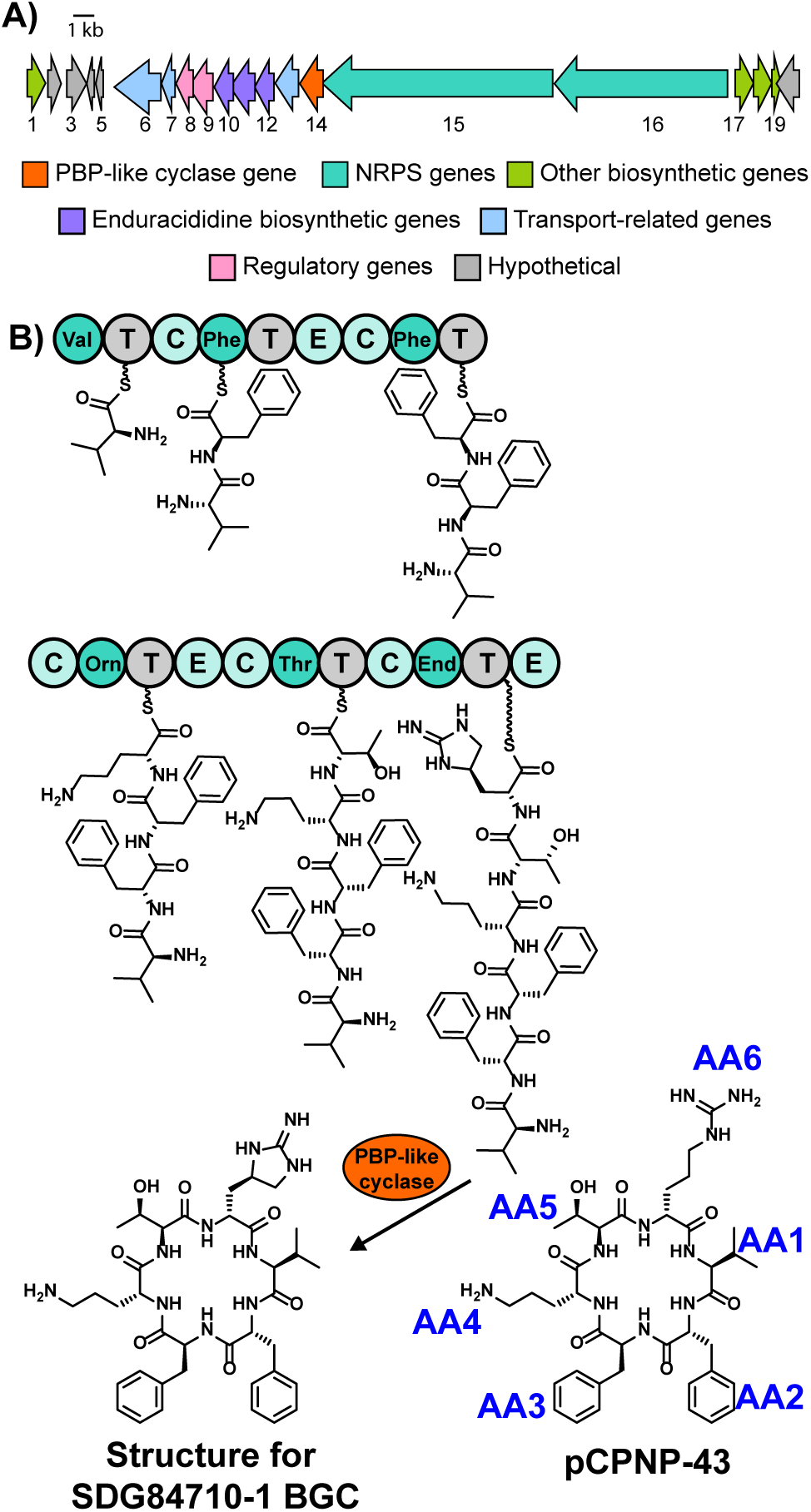
BGC for pNP-43. A) NRPS BGC including the PBP-like cyclase SDG84710.1. B) NRPS modules and amino acid predictions by PRISM. AA# refer to the amino acid position of pNP-43.

Due to its improved activity against the antibiotic resistant strain, we chose to study the mechanism of action of **pNP-43d**. Many cyclic peptides, including colistin (**Figure 5A**), are known to cause bacterial cell lysis. Specifically, colistin is known to interact with Lipid A via its 5 positively charged amino acids, displace divalent cations, and weaken the bacterial outer membrane of Gram-negative bacteria.^50^ This ultimately allows the peptide to enter the cell, where its additional activities have been postulated to cause cell death. Due to the positively charged amino acids on **pNP-43d**, we chose to test it for its ability to lyse bacterial cells using a previously reported Sytox green assay.^51^ **pNP-43d** clearly resulted in bacterial cell lysis at concentrations varying from 4 to 32 times the MIC for both wild type and antibiotic resistant *A. baumannii* (**Figure 5B** and **Figure S7**). Others have observed that colistin does not induce cell lysis at its MIC and suggested that while its lytic abilities likely contribute to its activity, it has other major mechanisms of action.^52^ Based on our results here, similar conclusions can be drawn for **pNP-43d**.

**Figure 5.**
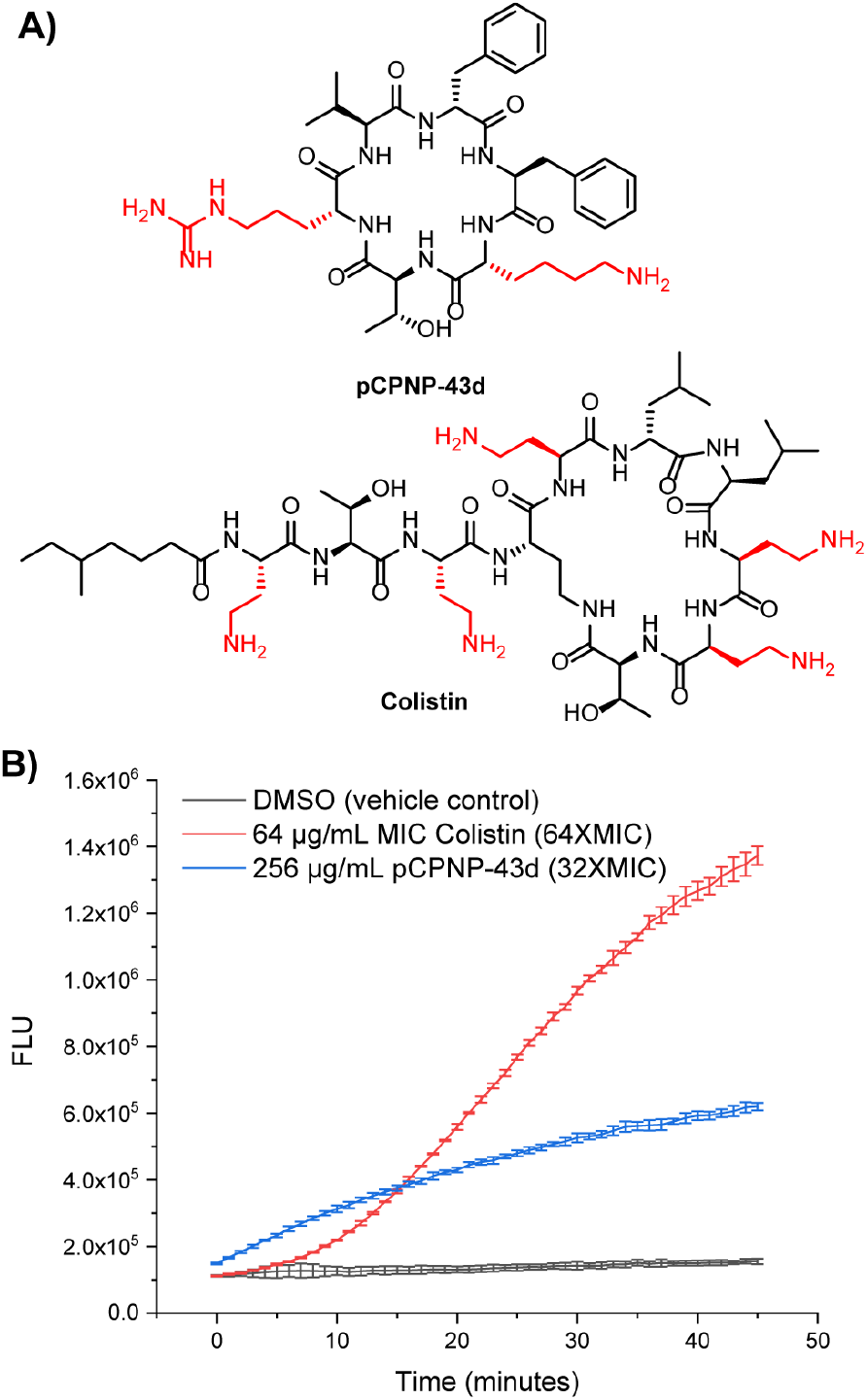
Mechanism of action studies. A) Chemical structures of pNP-43d and colistin with basic residues indicated in red. B) Representative data from Sytox Green lysis assay with *A. baumannii* 19606. Error bars are standard deviation from 3 technical replicates. *N* = 3

## CONCLUSIONS

Described herein is the development of SNaPP, a method to greatly expedite the discovery of bioactive molecules inspired by NPs. Cyclic peptides were chosen as an initial target due to their history as important sources of medicines along with the established bioinformatics approaches for predicting the peptide sequences. Head-to-tail peptides were targeted by identifying NRPS BGCs that co-occur with genes from a recently discovered family of stand-alone cyclases, the PBP-like cyclases. This approach allowed for identification of 131 unique and novel cyclic peptides. 51 diverse pNPs were chemically synthesized and tested for antibiotic activity. Approximately 30% of pNPs had activity with several showing very promising activity against difficult-to-treat Gram-negative bacteria. As prediction softwares for NP BGCs improve, this strategy will only increase in its utility. Overall, SNaPP is a powerful method for the rapid identification of biologically inspired lead molecules.

## ASSOCIATED CONTENT

### Author Contributions

M.A.H. and E.I.P. designed the experiments. M.A.H., C.S., I.W., A.F., J.G., and E.I.P. performed the bioinformatics predictions. M.A.H., C.S., S.N., Z.B., and B.B synthesized the pNPs. E.I.P. performed the antibiotic and hemolysis screening. R.M. performed the mammalian cell screening. M.A.H. and E.I.P. wrote the paper. S.N. edited the paper.

### Funding Sources

1R35GM138002-01 to E.I.P.

Frederick N. Andrews Fellowship to S.N.

P30 CA023168 to the Purdue Center for Cancer Research

## ACKNOWLEDGMENT

We are grateful to P.J. Hergenrother (Univ. of Illinois at Urbana-Champaign) and W.W. Metcalf (Univ. of Illinois at Urbana-Champaign) for providing the bacterial pathogens used in this study. Phillip Wyss provided advice and assistance on programming. This work was supported by a grant from the National Institutes of Health (1R35GM138002-01 to E.I.P.) and the Frederick N. Andrews Fellowship from the Department of Medicinal Chemistry and Molecular Pharmacology at Purdue University (to S.N.). The authors acknowledge the support from the Purdue Center for Cancer Research, NIH grant P30 CA023168.

## ABBREVIATIONS

NP: natural product
pNP: predicted natural product
BGC: biosynthetic gene cluster
SNaPP: <u>Bi</u>ologically <u>I</u>nspired <u>C</u>hemically <u>C</u>reated <u>L</u>eads
NRPS: non-ribosomal peptide synthetase
NRP: non-ribosomal peptide
PBP: penicillin-binding protein
SSN: sequence similarity network

