## Supplemental Materials, Figures, and Tables for "Synthetic Natural Product Inspired Peptides"

**Table of Contents**

**Materials and Methods……………………………………………………………………S3-S5**

**Figure S1: Comparison of PRISM and antiSMASH predictions………………………………S6**

**Figure S2: Tanimoto Similarity Tree……………………………………………………………….S7**

**Figure S3: Sequence Similarity Network for PBP-like cyclases……………………………S8**

**Figure S4: BiG-SCAPE network of PBP-like cyclase and NRPS containing BGCs……..S9**

**Figure S5: Overview of chemical synthesis of pNPs ..…………………………………..…S10**

**Figure S6: pNPs with antibiotic activity………………..…………………………………….S11**

**Figure S7: Bacterial Lysis ……………………………..……………………………………S12**

**Table S1: pNP Library……………………………………………………………………….S13-S14**

**Table S2: pNPs with activity against Gram-negative bacteria………………………………S15**

**Table S3: pNPs with activity against Gram-positive bacteria ……………………………S16**

**Table S4: Predictions for genes in the pNP-43 BGC……………………………………………S17**

**Table S5: Predictions for genes in the pNP-43 BGC……………………………………………S18**

### Materials and Methods

***General Information***. Solvents were purchased from Fisher Scientific and used without further purification. Fmoc amino acids, coupling reagents, were purchased from Chem-Impex International. 2-CTC resin was purchased from ChemPep Incorporated. All other reagents were purchased from commercially available sources (Sigma Aldrich, Acros Organics, Oakwood Chemical, TCI Chemicals), and used without further purification. See Key Resources Table for more information.

***Bacterial Strains***. All strains used in this study except the *Bacillus* strain were obtained from Professor Paul Hergenrother (UIUC). The *Bacillus* strain was obtained from Professor William Metcalf (UIUC). *E. coli* ATCC 25922 (wild type, WT) and BAA-2469 (resistant, R), *K. pneumonia* ATCC 27736 (WT) and BAA-2146 (R), *A. baumannii* ATCC 19606 (WT) and KB349 (R), and *P. aeruginosa* PAO1 (WT) and PA1000 (R) were grown on Mueller Hinton Broth 2 (Sigma Aldrich). *S. aureus* ATCC 29213 (WT) and NRS3 (R), *Enterococcus* species ATCC 19433 (WT) and S235 (R), and *B. subtilis* 6633 (WT) were maintained on Bacto Brain Heart Infusion.

***Prediction of cyclic peptide structure***. The accession numbers for the top 500 hits from the SurE BlastP were downloaded and used as the input for RODEO [38]. Biosynthetic gene clusters were then manually analyzed for the presence of non-ribosomal peptide synthetase (NRPS) genes. If an NRPS was at the end of a contig, the cluster was not considered further. If the NRPS was not at the end of the contig, the FASTA file for the cluster was then analyzed using both PRISM 4.0 [17] and antiSMASH 5.0 [16]. Generally, both programs agreed well. Initial structures were assigned based on the PRISM results (see Supplementary Excel Document). Derivatives were designed based on results from both programs.

***Tanimoto similarity analysis.*** Tanimoto similarity analysis was accomplished with ChemMine Tools [52] using the following parameters for hierarchical clustering: Display values: Z-scores; Linkage method: single; Heatmap: distance matrix.

***Sequence similarity analysis.*** Sequence similarity analysis of the PBP-like cyclases was accomplished using the EFI-Enzyme Similarity Tool [42] and visualized using Cytoscape 3.6.1 [53]. An alignment score of 120 was used for generating the networks in this paper.

***BiG-SCAPE analysis***. BiG-SCAPE analysis[43] was performed on the 316 BGCs containing both a PBP-like cyclase and an NRPS. The antiSMASH outputs from the prediction of the cyclic peptide structure were used an inputs for BiG-SCAPE. The output was visualized using Cytoscape 3.6.1.

***General Procedure for Resin Loading***. 2-Chlorotrityl chloride (2-CTC) resin (1.0 g, 0.77 mmol/g, 0.77 mmol, 100-200 mesh), was swelled in DMF for 30 min, drained, and treated with a solution of Fmoc-protected amino acid (2.3 mmol) and DIEA (537 µL, 3.09 mmol) in DMF (11 mL). The resulting mixture was gently agitated for 2 h, after which, the resin was filtered and washed with DMF (2 x 5 mL). Remaining unreacted Cl groups were capped by agitating the resin with 11 mL $CH_2Cl_2$-MeOH-DIEA (17:2:1) for 20 min. The resin was filtered and washed with $CH_2Cl_2$ (3 x 5 mL), MeOH (3 x 5 mL), and dried under vacuum for 1 h. The resin loading was determined by treating an aliquot (1-3 mg) of the dried resin with piperidine-DMF (1:4) and observing the UV absorbance of the piperidine-dibenzofulvene adduct at 301 nm ($\varepsilon$ = 7800 $M^{-1}$ $cm^{-1}$).

***General procedure for manual SPPS***. 5 mL fritted polypropylene syringes (Torviq) were used as reaction vessels for all manual SPPS and cleavage steps. Pre-loaded 2-CTC resin (0.05 mmol) was swelled in DMF for 30 min, drained, and treated with piperidine-DMF (1:4, 3 mL, 1 x 15 min).

The resin was filtered and washed with DMF (2 x 3 mL) then $CH_2Cl_2$ (2 x 3 mL). In a separate flask, DIC (31 µL, 0.2 mmol) was added to a solution of Fmoc-AA-OH (0.2 mmol) and Oxyma Pure (0.2 mmol) in DMF (1.7 mL). Following a 5 min preactivation period, the resulting solution was added to the resin and the mixture agitated for 1 h. The resin was filtered and washed with DMF (3 x 2 mL) then $CH_2Cl_2$ (3 x 2 mL) and Kaiser ninhydrin[54] test performed to determine reaction completion. Deprotection and coupling cycles were repeated until the desired peptide sequence was complete.

***General procedure for automated SPPS*.** Linear peptides were synthesized on the 0.05 mmol scale using a PS3 peptide synthesizer (Gyros Protein Technologies). DIC, Oxyma Pure, and Fmoc-AA-OH (6 equiv, 0.3 mmol each) were used to accomplish couplings in 1 h, and Fmoc removal was achieved using piperidine-DMF (1:4, 2 x 5 min). Pre-loaded 2-chlorotrityl chloride resin (prepared as described above) was used for all syntheses.

***General procedure for peptide cleavage.*** The peptide-linked resin was swelled in DMF (1 x 15 min), drained, and treated with 20% piperidine-DMF (1 x 15 min) to remove N-terminal Fmoc group. The resin was drained and washed with DMF (3 x 2 mL) then $CH_2Cl_2$ (3 x 2 mL) and a Kaiser ninhydrin[54] test was performed to verify successful deprotection. The resin was treated with a 3 mL of a mixture of HFIP-$CH_2Cl_2$ (1:4) for 30 min and the filtrate concentrated under reduced pressure. The resulting residue was taken up in ~5 mL 50% $H_2O$-$CH_3CN$, frozen, and lyophilized to afford crude, side chain-protected, linear peptides that were used without further purification.

***General procedure for peptide cyclization and global deprotection.*** To a solution of crude linear peptide (~0.05 mmol) and PyBop (78 mg, 0.15 mmol) in DMF (40 mL) was added DIEA (52 µL, 0.30 mmol). This solution was agitated overnight (17-24 h) and concentrated under reduced pressure. 10 mL of 50% $H_2O$-$CH_3CN$ was added to the residue, the mixture vortexed, then centrifuged to afford a precipitate which was isolated by removal of the supernatant. The resulting solids were washed with an additional 10 mL 50% $H_2O$-$CH_3CN$, centrifuged, and isolated as before. The solids were frozen and lyophilized to remove residual solvent. *Note: in rare cases where these conditions do not afford the cyclic peptides as precipitates, the crude peptides were purified at this stage by RP-HPLC ($CH_3CN/H_2O$ as the eluent). See Table S1 for more details.* The crude material was treated with 3 mL of a mixture of TFA-$CH_2Cl_2$-TIPS (50:45:5) for 2 h, volatiles removed by a stream of air, and peptide precipitated with 2 mL of MTBE. Solids were collected, washed once with MTBE, dissolved in $H_2O$-$CH_3CN$, frozen, and lyophilized to afford cyclic peptides that were generally >90% pure.

***HPLC Methods****.* HPLC analysis and purification was performed on an Agilent Technologies 1260 Infinity II preparative HPLC system using a 1260 variable wavelength detector (measuring at 214 nm and 254 nm). A Luna C18 reverse phase 5 µm, 150 x 4.6 mm column (Phenomenex) was used for purity analysis, and a Luna C18 reverse phase, 5 µm, 150 x 21.2 mm column (Phenomenex) was used for purification. Solvent A: water with 0.1% formic acid, solvent B: acetonitrile with 0.1% formic acid. For purity analysis, the following gradient was used: (A:B, 1 mL/min): 95:5, 0 min; 95:5, 1 min; 5:95, 20 min; 5:95, 25 min; 95:5, 30 min. For the preparatory HPLC runs, several different methods were employed. Method A2 (purification of linear peptide): (A:B, 20 mL/min): 95:5, 0 min; 95:5, 1 min; 5:95, 20 min; 5:95, 25 min; 95:5, 30 min. Method B2 (purification of cyclic peptide): (A:B, 20 mL/min): 95:5, 0 min; 95:5, 1 min; 5:95, 20 min; 5:95, 25 min; 95:5, 30 min. Method B3: ): (A:B, 20 mL/min): 95:5, 0 min; 95:5, 1 min; 60:40, 20 min; 5:95, 25 min; 95:5, 30 min. Further information about preparatory HPLC runs can be found in Table S1.

***Mass Spectrometry*****.** Mass spectra (MS) were recorded on an Advion Expression CMS single quadrupole mass spectrometer using electrospray ionization (ESI).

**Antibacterial activity analysis.** Antibacterial activity analysis for all bacteria was performed using the microdilution broth method as outlined by the Clinical and Laboratory Standards Institute (CLSI).[46] Mueller Hinton Broth 2 (MH, Sigma-Aldrich, 90922) was used for all testing. Testing was performed as previously described.[55] Turbidity (OD600) of the wells was determined using a SpectraMax iD3 platereader (Molecular Devices). For the compounds that hit during initial screens, minimum inhibitory concentrations were determined. A minimum of three biological replicates were performed. Ciprofloxacin was used as a control in these assays.

**Anticancer Testing.** A549 non-small cell lung cancer cells (ATCC CCL-185) were obtained directly from ATCC and used within 30 passages. A549 cells were maintained in RPMI 1640 medium supplemented with 10% fetal bovine serum, 100 U/mL penicillin, and 100 ug/mL streptomycin. For anticancer testing, cells were seeded at 2000 cells per well in 96 well plates and allowed to adhere overnight. Cells were then treated with compound at 16 ug/mL (1% DMSO final) or vehicle control for 48 hours. Viability was assessed using Alamar Blue. Specifically, resazurin (Sigma Aldrich, R7017) was dissolved at 440 µM in sterile PBS. 20 µL was then added to each well and incubated for 4-8 h at 37 °C. Fluorescence was measured using a SpectraMax iD3 platereader (Excitation: 550 nm, Emission 590 nm). Percent death was calculated by subtracting the background from all wells and setting 0% death to vehicle treated controls.

**Hemolysis Assay.** Hemolysis assays were based on a previously described method.[56] Human Whole Blood was purchased from BioIVT and used prior to its expiration date. 100 µL of blood was aliquoted into a 1.5 mL Eppendort tube and 500 µL of sterile 0.9% NaCl was added. Tubes were gently inverted to mix and then centrifuged at 500Xg for 7 minutes. Supernatant was carefully removed and the pellet was washed 2X with 500 µL of 0.9% NaCl. The pellet was then resuspended in 800 µL of Red Blood Cell (RBC) buffer (10 mM $Na_2HPO_4$, 150 mM NaCl, 1 mM $MgCl_2$, pH 7.4). To evaluate hemolytic activity of pNPs, 4 µL of a 1.6 mg/mL DMSO stock (54 µg/mL final working concentration) was transferred to a well of a 96 U-well plate. Negative control wells contained 4 µL of DMSO and positive controls contained 4 µL of 30% Triton X-100. To each well was then added 76 µL of RBC buffer and 40 µL of the resuspended red blood cells. This was incubated for 1h at 37 °C. The plate was then spun at 500Xg for 5 min and the supernatants from each sample (75 µL) were transferred to a flat-well 96 well plate. The absorbance of these supernatants at 540 nm was then measured using a SpectraMax iD3 platereader. Percent hemolysis was calculated relative to the average absorbance values for the positive and negative controls. A minimum of three biological replicates was performed.

**Bacterial Lysis Assay.** Bacterial lysis assays were based on a previously described method.[50] Briefly, 50 µL an overnight culture of *A. baumannii* was used to inoculate 5 mL of fresh MH medium. The culture was allowed to grow to mid-logarithmic phase (usually ~2 h). The bacteria was collected and washed 3 times with 5 mM HEPES (pH 7.4) supplemented with 20 mM glucose. After washing, bacteria were resuspended in 1 mL of 5 mM HEPES (pH 7.4) supplemented with 20 mM glucose and 100 mM KCl. *A. baumannii* suspensions of ~1E8 CFU were mixed with SYTOX Green (Invitrogen, 0.5 µM final concentration) and incubated for 15 minutes at room temperature in the dark. A 2X stock of compound or vehicle control was then mixed with the bacteria suspension and immediately transferred to a black clear bottom 96 well plate. Colistin was used as a positive control, and DMSO was used as a negative control. Bacterial cell lysis was monitored by the uptake of SYTOX green using a SpectraMax iD3 platereader (Excitation: 480 nm; Emission: 522 nm; read every 1 minute for 60 minutes).

**Figure S1: Comparison of PRISM and antiSMASH predictions.** A) Average Tanimoto value for predicted peptides from BGCs of 5 known natural products (NPs) from the dataset (mannopeptimycin, ulleugymycin, desoatmide, surugamide I, and surugamide F) compared to actual structure of the natural product (NP) along with the standard deviation. No significant different was observed between the Tanimoto values of the structures predicted from PRISM versus antiSMASH compared to the true structures (Student's T-test). B) Average Tanimoto value for peptides predicted by PRISM versus antiSMASH (blue) compared to Tanimoto value of PRISM peptides compared to a ring of glycines with the same number of amino acids (orange). (***p < 0.001, **p < 0.01).

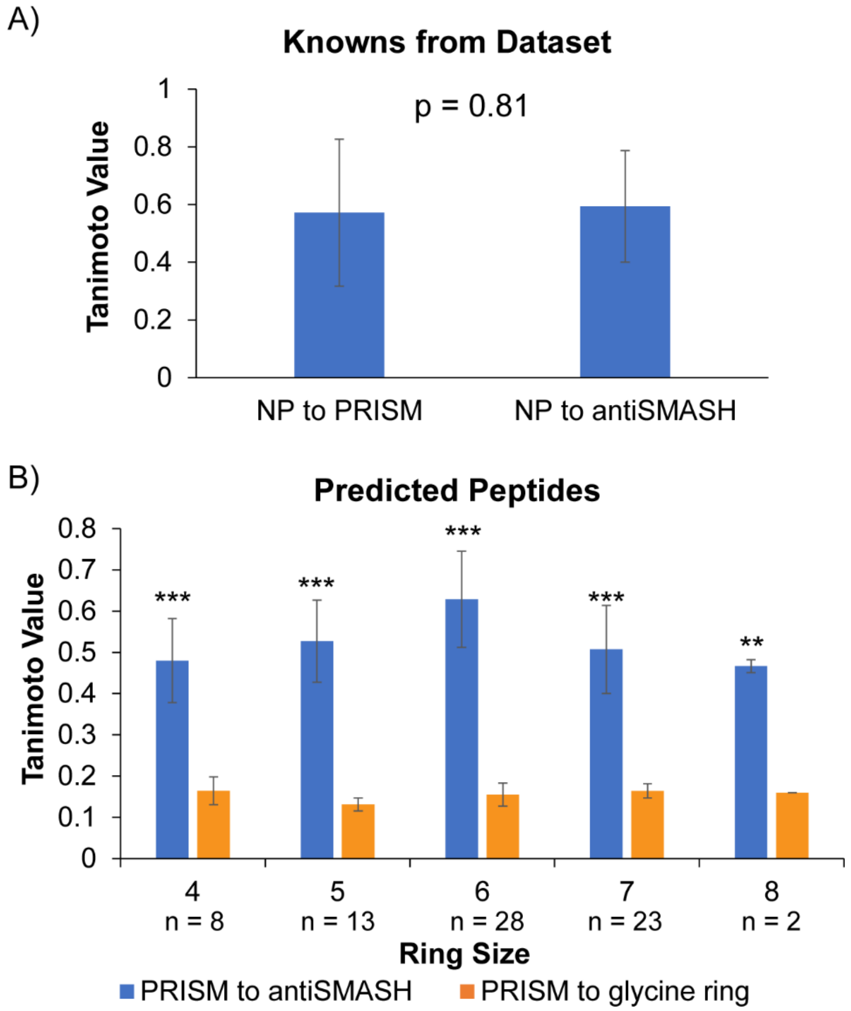

**Figure S2: Overview of pNP Library.** A) Tanimoto tree with synthesized pNPs indicated. Green circles are pNPs with the exact structure predicted. Blue squares are pNPs with slight modifications (*e.g.* Phg substituted for Hpg). B) pNPs distribution with the number of unique and novel cyclic peptides noted in light green and the number of synthesized pNPs indicated in dark green. Note the number of synthesized pNPs is also indicated on the graph.

**A)**

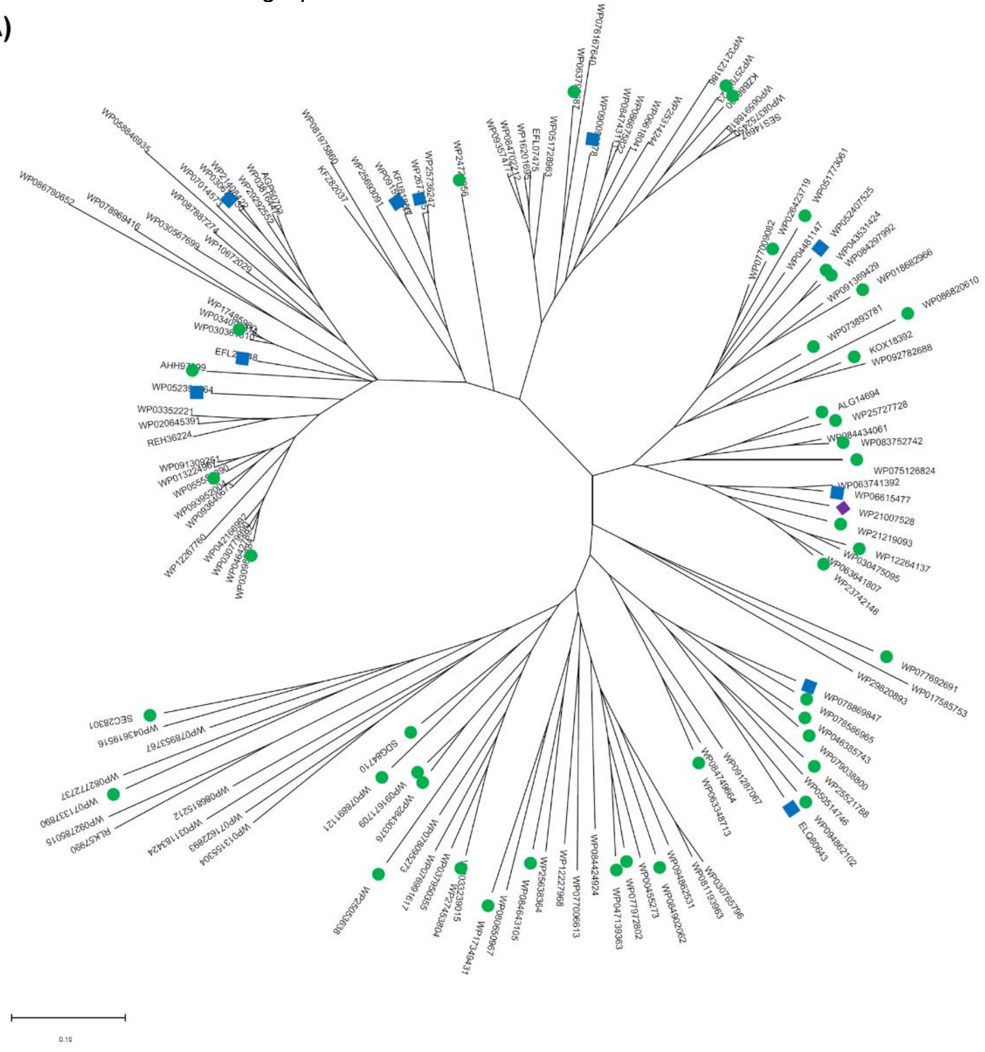

**B)**

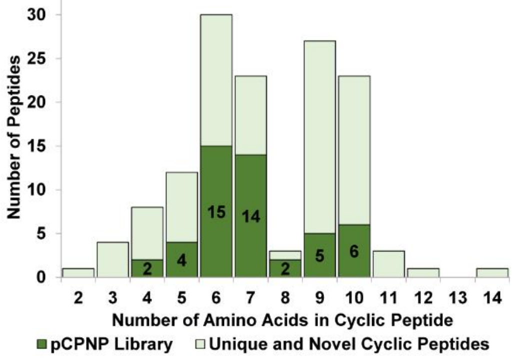

**Figure S3: Sequence Similarity Network for PBP-like cyclases**. The size (number of amino acids) of the predicted cyclic peptide product is indicated by the color of the nodes. Nodes with thick red outlines have been synthesized.

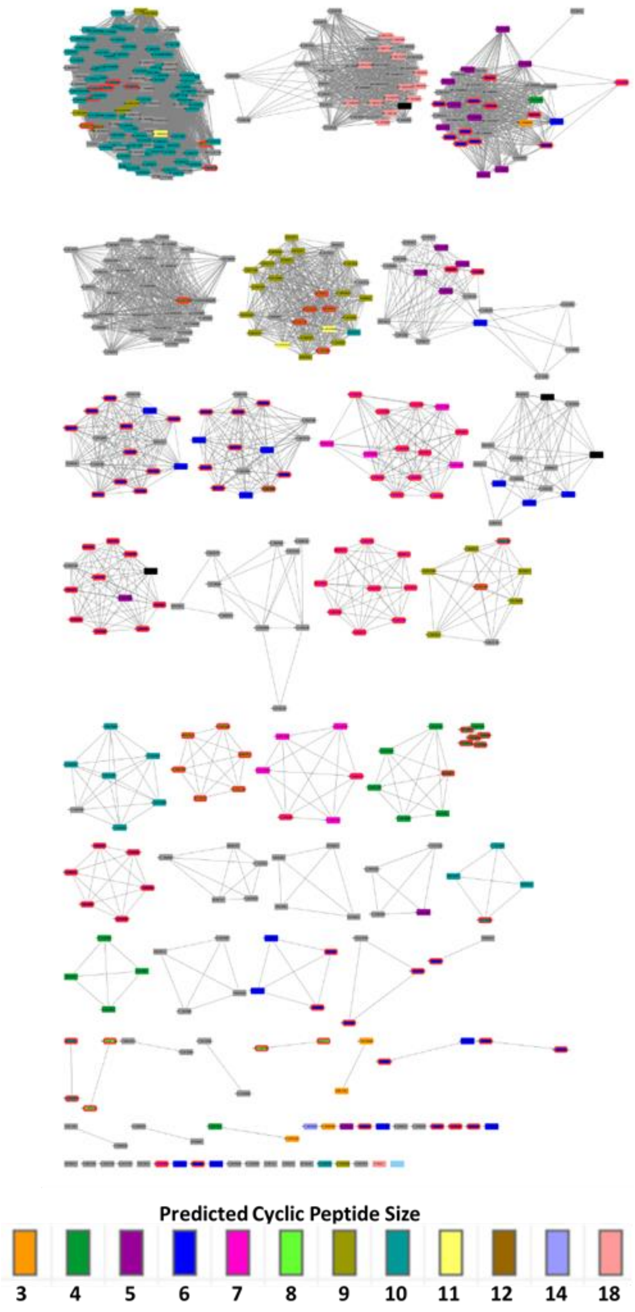


E = -120

**Figure S4: BiG-SCAPE network of PBP-like cyclase and NRPS containing BGCs.** Each circle represents a family (closely related) of BGCs. Branches to other circles indicate clans (more distantly related BGCs). The size (number of amino acids) of the predicted cyclic peptide product is indicated by the color of the nodes. Nodes with thick red outlines have been synthesized.

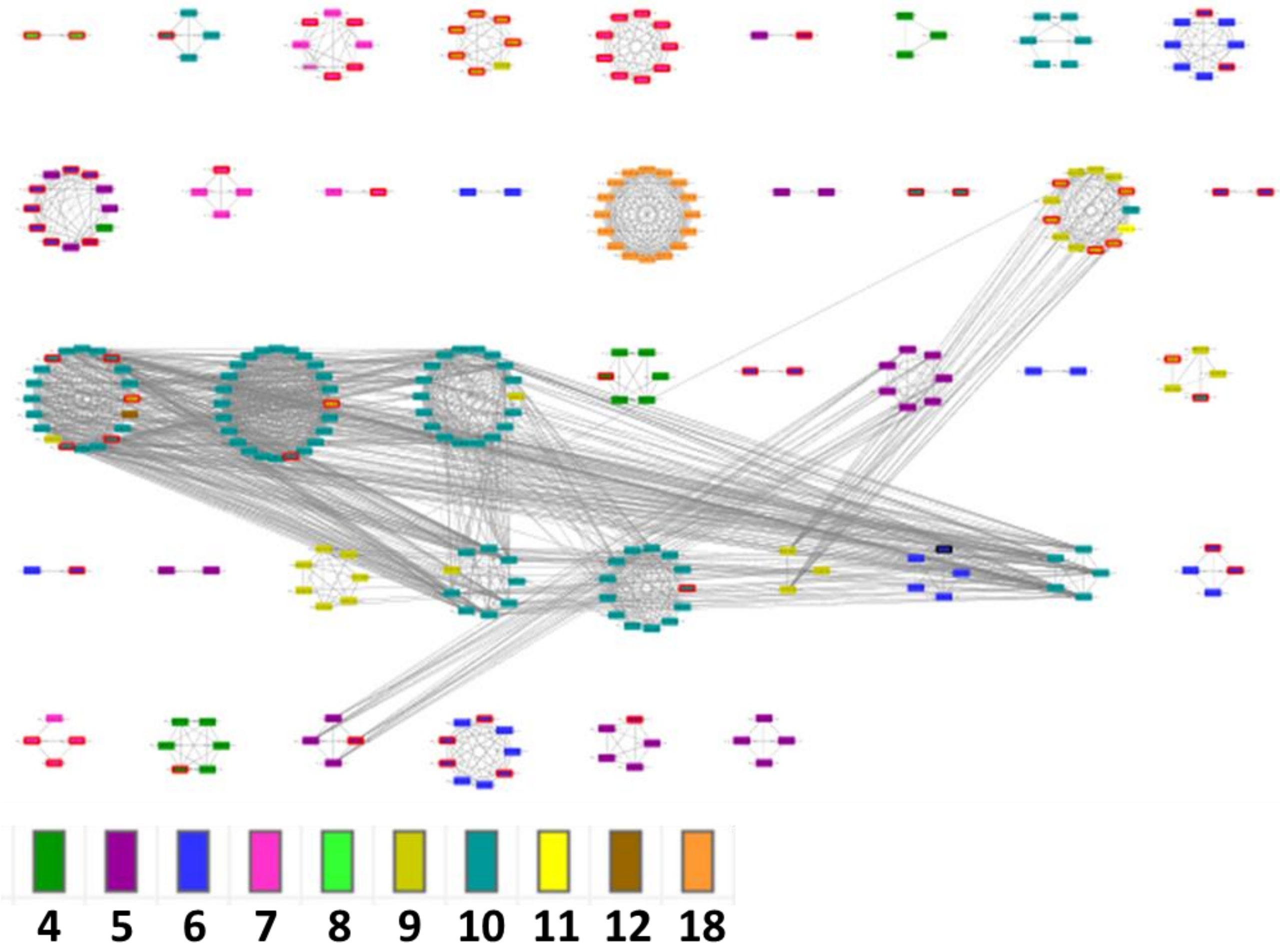

**Figure S5: Overview of chemical synthesis of pNPs.** Cyclic peptides were synthesized using Fmoc / *t*Bu solid-phase peptide synthesis (SPPS) on the 2-chlorotrityl chloride resin (2-CTC). Fmoc removal was achieved using 20% piperidine in DMF and coupling was promoted using Oxyma and DIC. Cycles of coupling and deprotection were repeated until desired sequence was completed. HFIP was used to release the linear peptide from the resin while leaving side chain protecting groups intact. Following a PyBop-promoted solution-phase cyclization, global deprotection was achieved using TFA to afford desired cyclic peptides. Often, this process afforded peptides that were used without further purification. However, in some instances HPLC purification at Step A, Step B, or both were required (see Table S1).

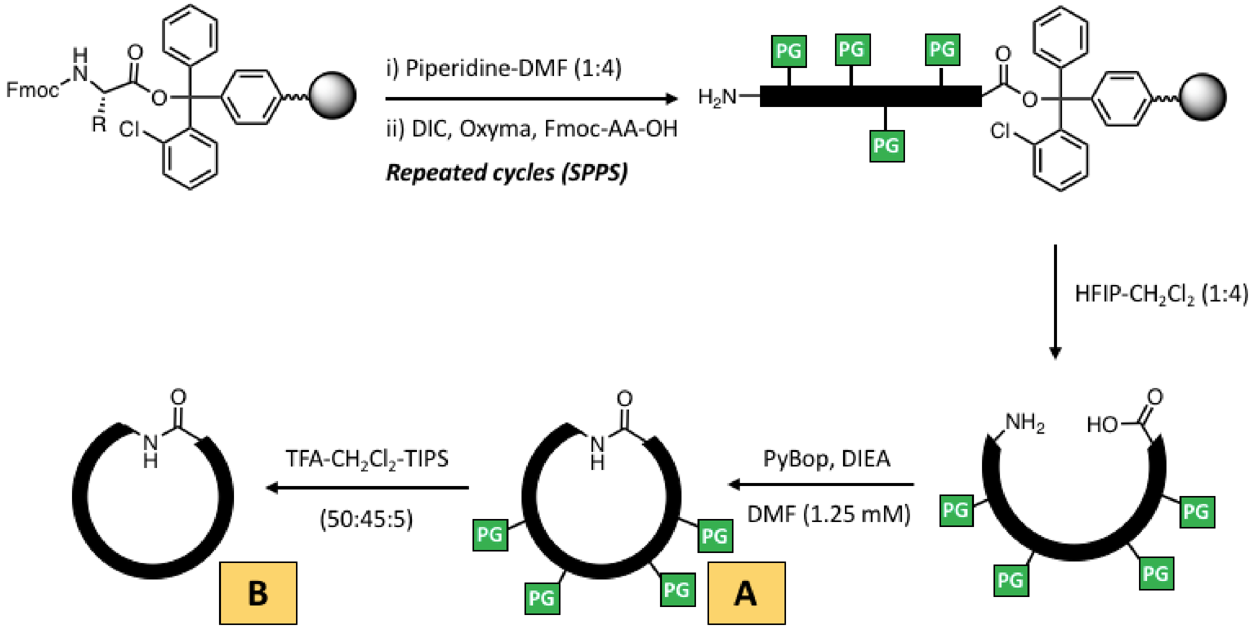

**Figure S6: pNPs active against Gram-positive bacteria.** A) Structures of pNPs that hit against Gram-positive bacteria. B) Structures of **pNP-43** derivatives.

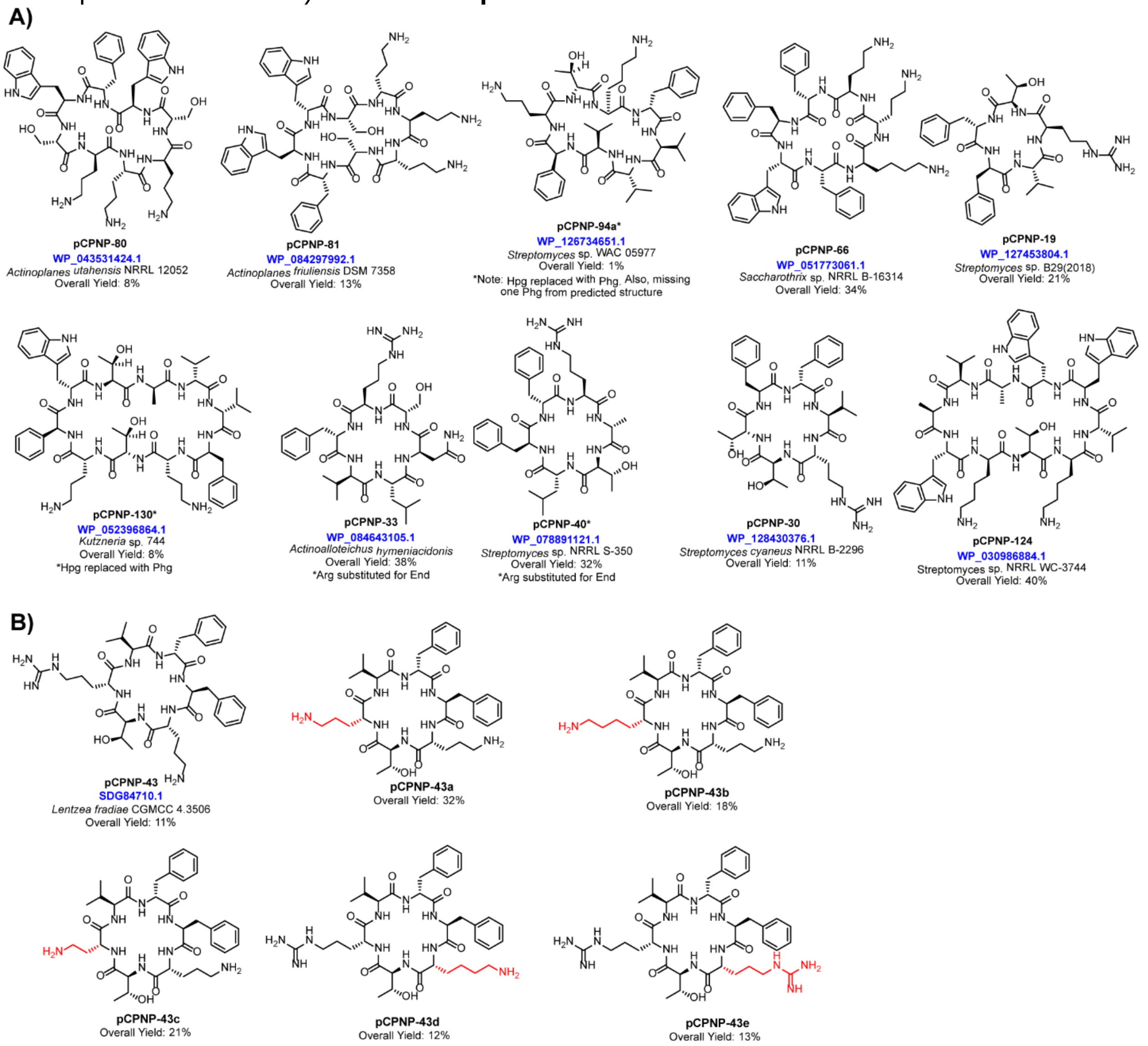

**Figure S7: Bacterial Lysis.** A) Representative data from Sytox Green lysis assay with *A. baumannii* KB349. B) Representative data from Sytox Green lysis assay with *A. baumannii* 19606 treated with different concentrations of colistin. C) Representative data from Sytox Green lysis assay with *A. baumannii* 19606 treated with different concentrations of **pNP-43d**. Error bars are standard deviation from 3 technical replicates. *N* = 3

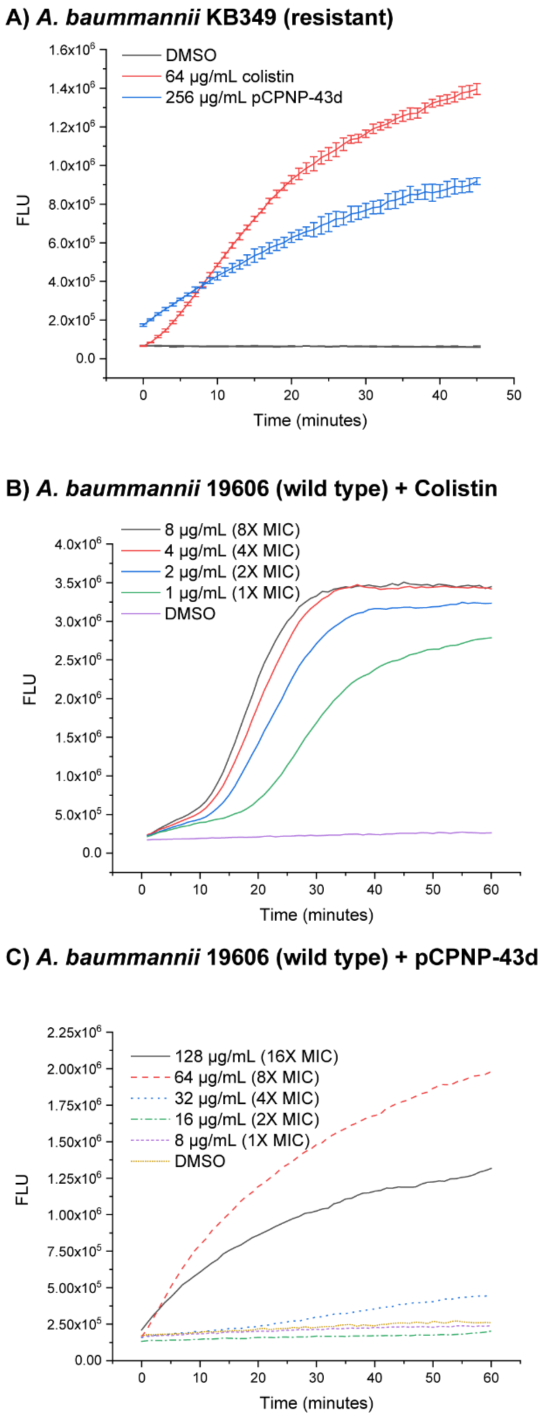

### Table S1: pNP Library

| #AA | pNP Identifier | Accession no. | Structure | $[M+H]^+$ (formula, calcd) | Isolated yield | | Purification* |
|---|---|---|---|---|---|---|---|
| | | | | | % | mg | |
| 4 | pNP-11 | WP_071337890.1 | *cyclo*[Val-Phe-D-Orn-Gly] | 418.1 ($C_{21}H_{31}N_5O_4$, 418.2) | 3 | 1 | A1, B** |
| 4 | pNP-12 | SEC28301.1 | *cyclo*[D-Arg-D-Ser-Val-Phe] | 490.3 ($C_{23}H_{36}N_7O_5$, 490.3) | 4 | 1.4 | A1, B** |
| 5 | pNP-16 | WP_077692691.1 | *Cyclo*[Phe-Val-D-Phg-Trp-D-Lys] | 693.36 ($C_{39}H_{47}N_7O_5$, 694.0) | 18 | 6.2 | A1, B2 |
| 5 | pNP-19 | WP_127453804.1 | *cyclo*[D-Phe-Val-D-Arg-D-Thr-Phe] | 651.5 ($C_{33}H_{47}N_8O_6$, 651.4) | 21 | 8.0 | A1, B1 |
| 5 | pNP-23 | WP_125053638.1 | *cyclo*[Phe-D-Phe-D-Leu‡-Thr-Gly] | 566.7 ($C_{30}H_{40}N_5O_6$, 566.3) | 42 | 12 | A1, B1 |
| 5 | pNP-28 | WP_079165356.1 | *cyclo*[Leu-D-Orn-Thr-D-Leu-D-Phe] | 589.8 ($C_{30}H_{49}N_6O_6$, 589.4) | 3 | 0.8 | A1, B2 |
| 6 | pNP-30 | WP_128430376.1 | *cyclo*[Val-D-Arg-Thr-D-Thr-Phe-D-Phe] | 752.4 ($C_{37}H_{54}N_9O_8$, 752.4) | 11 | 4.6 | A1, B1 |
| 6 | pNP-33 | WP_084643105.1 | *cyclo*[D-Val-Leu-D-Asn-Ser-D-Arg‡-Phe] | 717.6 ($C_{33}H_{53}N_{10}O_8$, 717.4) | 38 | 16 | A1, B1 |
| 6 | pNP-34 | WP_046385743.1 | *cyclo*[Gly-Phe-D-Val-D-Arg-Lys-D-Phe] | 735.4 ($C_{37}H_{55}N_{10}O_6$, 735.4) | 10 | 5 | A2, B1 |
| 6 | pNP-35 | WP_094862102.1 | *cyclo*[Leu-D-Val-D-Orn-Lys-D-Phe-Gly] | 659.4 ($C_{33}H_{55}N_8O_6$, 659.5) | 11 | 4.2 | A1, B2 |
| 6 | pNP-36 | WP_077972802.1 | *cyclo*[Arg-D-Thr-Asn-D-Val-Val-D-Phe] | 716.4 ($C_{33}H_{52}N_{10}O_8$, 717.4) | 79 | 28 | A1, B1 |
| 6 | pNP-37 | WP_079038800.1 | *cyclo*[Phe-D-Val-D-Orn-Lys-D-Phe-Gly] | 693.4 ($C_{36}H_{52}N_8O_6$, 692.40) | 28 | 13 | A1,B1 |
| 6 | pNP-40 | WP_078891121.1 | *cyclo*[D-Phe-Arg‡-D-Ala-Thr-D-Leu-Phe] | 736.6 ($C_{37}H_{54}N_9O_7$, 736.4) | 32 | 14 | A1, B1 |
| 6 | pNP-43 | SDG84710.1 | *cyclo*[Phe-D-Orn-Thr-D-Arg‡-Val-D-Phe] | 765.5 ($C_{38}H_{57}N_{10}O_7$, 765.4) | 11 | 4.9 | A1, B3 |
| 6 | pNP-43a | SDG84710.1 derivative | *cyclo*[Phe-D-Orn-Thr-D-Orn-Val-D-Phe] | 723.4 ($C_{37}H_{55}N_8O_7$, 723.4) | 32 | 15 | A1, B1 |
| 6 | pNP-43b | SDG84710.1 derivative | *cyclo*[Phe-D-Orn-Thr-D-Lys-Val-D-Phe] | 737.4 ($C_{38}H_{57}N_8O_7$, 737.4) | 18 | 8.6 | A1, B1 |
| 6 | pNP-43c | SDG84710.1 derivative | *cyclo*[Phe-D-Orn-Thr-D-Dab-Val-D-Phe] | 709.8 ($C_{36}H_{53}N_8O_7$, 709.4) | 21 | 9.7 | A1, B1 |
| 6 | pNP-43d | SDG84710.1 derivative | *cyclo*[Phe-D-Lys-Thr-D-Arg-Val-D-Phe] | 780.0 ($C_{39}H_{59}N_{10}O_7$, 779.5) | 12 | 5.9 | A1, B1 |
| 6 | pNP-43e | SDG84710.1 derivative | *cyclo*[Phe-D-Arg-Thr-D-Arg-Val-D-Phe] | 807.5 ($C_{39}H_{59}N_{12}O_7$, 807.5) | 13 | 6.9 | A1, B1 |
| 6 | pNP-44 | WP_047139363.1 | *cyclo*[Arg-D-Thr-Val-Asn-D-Val-D-Phe] | 716.4 ($C_{33}H_{52}N_{10}O_8$, 717.0) | 8 | 2.7 | A2, B1 |
| 6 | pNP-45 | WP_125521788.1 | *cyclo*[Arg-D-Val-D-Orn-Lys-D-Trp-Gly] | 741.7 ($C_{35}H_{57}N_{12}O_6$, 741.4) | 23 | 13 | A2, B1 |
| 6 | pNP-47 | WP_078869847.1 | *cyclo*[Trp-D-Val-D-Orn-Lys-D-Trp-D-Phe‡] | 861.7 ($C_{47}H_{61}N_{10}O_6$, 861.5) | 28 | 15 | A1, B1 |
| 6 | pNP-48 | WP_063348713.1 | *cyclo*[D-Val-Asn-D-Orn-Arg-D-Trp-Phe] | 817.9 ($C_{40}H_{57}N_{12}O_7$, 817.4) | 16 | 8.3 | A1, B3 |
| 6 | pNP-51 | WP_086820610.1 | *cyclo*[D-Orn-Orn-D-Orn-Phe-D-Phe-Phe] | 784.5 ($C_{42}H_{58}N_9O_6$, 784.4) | 15 | 8.4 | A1, B1 |
| 6 | pNP-53 | WP_078586965.1 | *cyclo*[Phe-D-Val-D-Orn-Lys-D-Trp-Gly] | 732.2 ($C_{38}H_{54}N_9O_6$, 732.4) | 9 | 4 | A1, B3 |
| 6 | pNP-56 | WP_052407525.1 | *cyclo*[Trp-D-Lys-Lys-D-Orn-Trp-Gly‡] | 800.7 ($C_{41}H_{58}N_{11}O_6$, 800.5) | 33 | 19 | A1, B1 |
| 7 | pNP-57 | ALG14694.1 | *cyclo*[D-Asp-Orn-D-Lys-Val-D-Phe-Trp-Ala] | 861.6 ($C_{43}H_{60}N_{10}O_2$, 861.0) | 29 | 16 | A1, B1 |
| 7 | pNP-58 | WP_125727728.1 | *cyclo*[D-Asp-Orn-D-Lys-Val-D-Trp-Trp-Ala] | 900.3 ($C_{45}H_{62}N_{11}O_9$, 900.5) | 36 | 20 | A1, B1 |
| 7 | pNP-59 | KOX18392.1 | *cyclo*[D-Phe-Phe-Thr-D-Lys-Lys-D-Orn-Phe] | 913.5 ($C_{48}H_{68}N_{10}O_8$, 913.1) | 41 | 25 | A1, B1 |
| 7 | pNP-60 | WP_083752742.1 | *cyclo*[D-Orn-Lys-D-Lys-Val-D-Phe-Trp-Ala] | 874.5 ($C_{45}H_{67}N_{11}O_7$, 874.1) | 29 | 17 | A1, B1 |
| 7 | pNP-61 | WP_121219093.1 | *cyclo*[Thr-D-Orn-Arg-D-Orn-Thr-D-Phe-Phe] | 881.6 ($C_{42}H_{65}N_{12}O_9$, 881.5) | 35 | 21 | A2, B1 |
| 7 | pNP-62 | WP_084434061.1 | *cyclo*[D-Orn-Orn-D-Lys-Val-D-Phe-Trp-Ala] | 860.5 ($C_{44}H_{65}N_{11}O_7$, 860.0) | 33 | 20 | A1, B1 |
| 7 | pNP-63 | WP_018682966.1 | *cyclo*[D-Trp-Trp-Ser-D-Lys-Lys-D-Lys-Phe] | 991.4, ($C_{52}H_{70}N_{12}O_8$, 991.2) | 13 | 8.8 | A1, B3 |
| 7 | pNP-64 | WP_026423719.1 | *cyclo*[Ser-D-Lys-Orn-D-Orn-Trp-D-Phe-Phe] | 924.6 ($C_{48}H_{66}N_{11}O_8$, 924.5) | 37 | 24 | A1, B1 |

| | | | | | | | |
|---|---|---|---|---|---|---|---|
| 7 | pNP-66 | WP_051773061.1 | *cyclo*[Trp-Phe-D-Lys-Orn-D-Orn-Phe-D-Phe] | 984.6 ($C_{54}H_{70}N_{11}O_7$, 984.5) | 34 | 22 | A1, B1 |
| 7 | pNP-69 | WP_073893781.1 | *cyclo*[D-Phe-Phe-Thr-D-Orn-Val-D-Orn-Phe] | 870.6 ($C_{46}H_{64}N_9O_8$, 870.5) | 36 | 20 | A1, B1 |
| 7 | pNP-70 | WP_075126824.1 | *Cyclo*[Ser-Trp-D-Trp-Thr-D-Lys-Orn-D-Orn] | 916.5 ($C_{45}H_{64}N_{12}O_9$, 917.1) | 2 | 0.8 | A1, B2 |
| 7 | pNP-72 | WP_084902062.1 | *cyclo*[D-Arg-Ser-D-Thr-D-Val-Thr-Phe-Gly] | 749.5 ($C_{33}H_{53}N_{10}O_{10}$, 749.4) | 4 | 2 | A2, B1 |
| 7 | pNP-76 | WP_106615477.1 | *cyclo*[Thr-D-Lys-Val-D-Orn-Thr-D-Phe-Phe] | 838.4 ($C_{42}H_{63}N_9O_9$, 838.5) | 12 | 6.6 | A1, B1 |
| 6 | pNP-76a | WP_106615477.1 derivative | *cyclo*[Thr-D-Lys-Val-Thr-D-Phe-Phe] | 724.6 ($C_{37}H_{53}N_7O_8$, 723.8) | 15 | 6 | A2, B1 |
| 7 | pNP-77 | WP_11264137.1 | *Cyclo*[Phe-D-Phe-Thr-D-Orn-Orn-D-Orn-Ser] | 824.5 ($C_{40}H_{60}N_{10}O_9$, 825.3 | 13 | 5.4 | A1, B1 |
| 7 | pNP-78 | WP_121007528.1 | *cyclo*[Thr-D-Orn-Asp-D-Lys-Thr-D-Phe-Phe] | 853.9 ($C_{41}H_{59}N_9O_{11}$, 853.4) | 90 | 43 | A1, B1 |
| 6 | pNP-78a | WP_121007528.1 derivative | *cyclo*[Thr-Asp-D-Lys-Thr-D-Phe-Phe] | 740.4 ($C_{36}H_{49}N_7O_{10}$, 739.8) | 27 | 13 | A2, B1 |
| 7 | pNP-79 | WP_123742148.1 | *cyclo*[Phe-Thr-D-Lys-Orn-D-Orn-Thr-D-Phe] | 853.0 ($C_{42}H_{64}N_{10}O_9$, 852.5) | 28 | 17 | A1, B1 |
| 8 | pNP-80 | WP_043531424.1 | *cyclo*[D-Trp-Ser-D-Orn-Orn-D-Orn-Ser-D-Trp-Phe] | 1036.6 ($C_{52}H_{70}N_{13}O_{10}$, 1036.5) | 8 | 5.2 | A1, B2 |
| 8 | pNP-81 | WP_124721056.1 | *cyclo*[Ala-D-Val-Val-Trp-Orn-D-Tyr-Trp-Ser] | 1006.4 ($C_{52}H_{67}N_{11}O_{10}$, 1006.2) | 13 | 7.2 | A1, B2 |
| 8 | pNP-88 | WP_078869847.1 | *cyclo*[Trp-D-Orn-Thr-D-Orn-D-Phe-Trp-D-Ala-D-Ala] ‡ | 991.7 ($C_{51}H_{67}N_{12}O_9$, 991.5) | 26 | 18 | A2, B1 |
| 8 | pNP-94 | WP_126734651.1 | *cyclo*[Val-D-Phg-D-Val-D-Val-Orn-D-Thr-Lys-D-Phe] | 921.0 ($C_{47}H_{72}N_{10}O_9$, 920.6) | 2 | 1.2 | |
| 9 | pNP-85 | WP_090096878.1 | *cyclo*[D-Asp-Phg‡-D-Val-D-Thr-Val-D-Val-Orn-D-Lys-Orn] | 1003.3 ($C_{47}H_{78}N_{12}O_{12}$, 1002.6) | 2 | 1.3 | A1, B4 |
| 9 | pNP-89 | WP_091596577.1 | *cyclo*[D-Trp-Val-D-Phg‡-D-Val-D-Val-Phg‡-Orn-D-Thr-Orn] | 1078.7 ($C_{56}H_{78}N_{12}O_{10}$, 1078.6) | 3 | 1.9 | A1, B1 |
| 9 | pNP-100 | WP_034091314.1 | *cyclo*[D-Val-Trp-D-Orn-Thr-D-Orn-Val-D-Trp-Trp-D-Ala] | 1155.6 ($C_{61}H_{82}N_{14}O_9$, 1156.6) | 22 | 15 | A1, B3 |
| 9 | pNP-101 | KZB86600.1 | *cyclo*[D-Asn-Val-D-Val-D-Val-D-Val-Val-Lys-D-Thr-Orn] | 953.3 ($C_{44}H_{80}N_{12}O_{11}$, 953.2) | 4 | 2.0 | A1, B1 |
| 10 | pNP-111 | AHH97299.1 | *cyclo*[D-Val-Thr-D-Orn-Orn-D-Orn-Trp-D-Ala-D-Val-D-Val-Phe] | 1144.8 ($C_{57}H_{88}N_{14}O_{11}$, 1144.7) | 7 | 3.8 | A1, B1 |
| 10 | pNP-112 | WP_063798687.1 | *cyclo*[D-Asn-Val-D-Val-D-Thr-Val-D-Val-Thr-Orn-D-Orn-Phe] | 1088.6 ($C_{51}H_{86}N_{13}O_{13}$, 1088.6) | 12 | 7.8 | A1, B2 |
| 10 | pNP-115 | WP_055551390.1 | *cyclo*[D-Orn-Thr-D-Orn-Val-D-Trp-Trp-D-Ala-D-Val-D-Ala-Phe] | 1189.7 ($C_{61}H_{85}N_{14}O_{11}$, 1189.6) | 35 | 25 | A1, B1 |
| 9 | pNP-123 | WP_03067330.1 | *cyclo*[D-Val-D-Trp-Trp-D-Orn-Thr-D-Orn-Val-D-Trp-Trp-D-Ala]‡ | 1157.1 ($C_{60}H_{80}N_{14}O_{10}$, 1156.6) | 40 | 31 | A1, B3 |
| 10 | pNP-124 | WP_030986884.1 | *cyclo*[Trp-D-Lys-Thr-D-Lys-Val-D-Trp-Trp-D-Ala-D-Val-D-Ala] | 1256.9 ($C_{65}H_{90}N_{15}O_{11}$, 1256.7) | 40 | 29 | A1, B1 |
| 10 | pNP-130 | WP_052396864.1 | *cyclo*[D-Orn-Thr-D-Orn-Phg‡-D-Trp-Thr-D-Ala-D-Val-D-Val-Phe] | 1165.3 ($C_{59}H_{83}N_{13}O_{12}$, 1165.6) | 7 | 4.8 | A1, B3 |

*See Figure S5 for more details. "A" refers to purification after release from the resin and "B" refers to purification after global deprotection. A1 = solids crashed out in 10 mL of 50% ACN and 50% $H_2O$; A2 = preparatory HPLC with the following gradient 0-1 min at 5% ACN (95% $H_2O$, 0.1% formic acid), 1-20 min gradient from 5-95% ACN, 20-25 at 95% ACN, and 25-30 at 5% ACN; B1 = solids crashed out of MTBE with no further purification needed (>90% pure); B2 = preparatory HPLC with the following gradient 0-1 min at 5% ACN (95% $H_2O$, 0.1% formic acid), 1-20 min gradient from 5-95% ACN, 20-25 at 95% ACN, and 25-30 at 5% ACN; B3 = preparatory HPLC with the following gradient 0-1 min at 5% ACN (95% $H_2O$, 0.1% formic acid), 1-20 min gradient from 5-40% ACN, 20-25 at 95% ACN, and 25-30 at 5% ACN; B4 = crashed out of 50% ACN and 50% $H_2O$

**Yields were too low to perform HPLC purification. Final compound was ~75% pure.

‡Derivative with substitution at this position for either enduracididine or hydroxyphenylglycine or derivative due to miscoupling. See the Supplementary Excel File for more details.

### Table S2: pNPs with activity against Gram-negative bacteria

| | *E. coli* | | *K. pneumoniae* | | *A. baumannii* | | *P. aeruginosa* | | Hemolysis | A549 toxicity |
|---|---|---|---|---|---|---|---|---|---|---|
| **Compound** | **WT** | **R** | **WT** | **R** | **WT** | **R** | **WT** | **R** | **53 μg/mL** | **16 μg/mL** |
| **pNP-23** | 16 (28) | >32 (>57) | >32 (>57) | >32 (>57) | >32 (>57) | >32 (>57) | >32 (>57) | >32 (>57) | <10% | <50% death |
| **pNP-43** | 32 (42) | 32 (42) | 32 (42) | >32 (>42) | 16 (21) | 32 (42) | >32 (>42) | >32 (>42) | <10% | <50% death |
| **pNP-43a** | 32 (44) | 32 (44) | >32 (>44) | >32 (>44) | >32 (>44) | >32 (>44) | >32 (>44) | >32 (>44) | <10% | <50% death |
| **pNP-43b** | 32 (43) | 32 (43) | 32 (43) | >32 (>43) | 16-32 (22-43) | 32 (43) | >32 (>43) | >32 (>43) | <10% | <50% death |
| **pNP-43c** | 32 (45) | 32 (45) | >32 (>45) | >32 (>45) | >32 (>45) | >32 (>45) | >32 (>45) | >32 (>45) | <10% | <50% death |
| **pNP-43d** | 32 (41) | 32 (41) | 32 (41) | >32 (>41) | 8-16 (10-21) | 8-16 (10-21) | >32 (>41) | >32 (>41) | <10% | <50% death |
| **pNP-43e** | 32 (40) | 32 (40) | 32 (40) | >32 (>40) | 16-32 (20-40) | 16-32 (20-40) | >32 (>40) | >32 (>40) | <10% | <50% death |
| **pNP-80** | 32 (31) | 32 (31) | 32 (31) | >32 (>31) | 16 (15) | 16-32 (15-31) | >32 (>31) | >32 (>31) | >10% | <50% death |
| **pNP-51** | >32 (>41) | >32 (>41) | >32 (>41) | >32 (>41) | >32 (>41) | >32 (>41) | 32 (41) | 16 (20) | <10% | <50% death |
| **pNP-111** | 32 (28) | 32 (28) | >32 (>28) | 32 (28) | >32 (>28) | 32 (28) | >32 (>28) | >32 (>28) | <10% | <50% death |
| **Cipro** | <0.1 (<1.4) | >32 (>87) | <0.1 (<1.4) | >32 (>87) | 1 (2.7) | >32 (>87) | 0.5-1.0 (1.4-2.7) | >32 | ND | ND |

WT: wild type; R: antibiotic resistant; Cipro: ciprofloxacin; WT *E. coli* : ATCC 25922; R *E. coli* : ATCC BAA-2469; WT *K. pneumoniae* : ATCC 27736; R *K. pneumoniae* : ATCC BAA-21469; WT *A. baumannii* : ATCC 19606; R *A. baumannii* : KB349; WT *P. aeruginosa* : PAO1; R *P. aeruginosa:* PA1000;

**Table S3: pNPs with activity against Gram-positive bacteria**

| Compound | *S. aureus* | | *Enterococcus* sp. | | *B. subtilis* | Hemolysis | A549 toxicity |
|---|---|---|---|---|---|---|---|
| | WT | R | WT | R | WT | 53 µg/mL | 16 µg/mL |
| **pNP-80** | 16 (15) | 16 (15) | 8-16 (8-15) | 8 (8) | 8 (8) | >10% | <50% death |
| **pNP-81** | 32 (31) | 8 (8) | 32 (31) | 8-16 (8-15) | 8 (8) | >10% | <50% death |
| **pNP-94a** | 32 (30) | 16 (15) | >32 (>30) | 16 (15) | 32 (30) | <10% | <50% death |
| **pNP-66** | >32 (>33) | 8 (8) | >32 (>33) | >32 (>33) | 16 (16) | <10% | <50% death |
| **pNP-130** | >32 (>27) | 16 (14) | >32 (>27) | >32 (>27) | >32 (>27) | >10% | <50% death |
| **pNP-40** | >32 (>43) | >32 (>43) | >32 (>43) | 4 (5) | >32 (>43) | >10% | <50% death |
| **pNP-33** | >32 (>45) | >32 (>45) | >32 (>45) | 32 (45) | >32 (>45) | <10% | <50% death |
| **pNP-30** | >32 (>43) | >32 (>43) | >32 (>43) | 4-8 (5-11) | >32 (>43) | >10% | <50% death |
| **pNP-19** | >32 (>49) | >32 (>49) | >32 (>49) | 8-16 (12-25) | >32 (>49) | >10% | <50% death |
| **pNP-124** | >32 (>25) | >32 (>25) | >32 (>25) | >32 (>25) | 16-32 (13-25) | <10% | <50% death |
| **Cipro** | <0.5 (<1.4) | >32 (>87) | 4-8 (11-22) | >32 (>87) | <0.5 (<1.4) | ND | ND |

WT: wild type; R: antibiotic resistant; Cipro: ciprofloxacin; WT *S. aureus* : ATCC 29213; R *S. aureus* : NRS3; WT *Enterococcus* : ATCC 19433; R *Enterococcus* : S235; WT *B. subtilis:* ATCC 6633

**Table S4: Predictions for genes in the pNP-43 BGC.** BlastP analysis was performed on 12-21-2020.

| Gene | Locus tag | Size (#AAs) | Proposed function |
|---|---|---|---|
| **1** | SDG84470.1 | 382 | glutathione-dependent formaldehyde dehydrogenase |
| **2** | SDG84473.1 | 189 | hypothetical protein |
| **3** | SDG84504.1 | 283 | GAF domain-containing protein |
| **4** | SDG84507.1 | 89 | hypothetical protein |
| **5** | SDG84537.1 | 103 | hypothetical protein |
| **6** | SDG84553.1 | 853 | ABC transport system permease protein |
| **7** | SDG84571.1 | 256 | ABC transporter ATP-binding protein |
| **8** | SDG84588.1 | 216 | response regulator transcription factor; two complenent transcriptional regulator, LuxR family |
| **9** | SDG84615.1 | 413 | signal transduction histidine kinase |
| **10** | SDG84621.1 | 272 | enduracididine biosynthesis enzyme MppR |
| **11** | SDG84651.1 | 382 | enduracididine biosynthesis enzyme MppQ |
| **12** | SDG84675.1 | 375 | enduracididine biosynthesis enzyme MppP |
| **13** | SDG84688.1 | 415 | MFS transporter |
| **14** | SDG84710.1 | 394 | Cubic0 group peptidase, beta-lactamase class C |
| **15** | SDG84734.1 | 4063 | Non-ribosomal peptide synthetase |
| **16** | SDG84748.1 | 3094 | Non-ribosomal peptide synthetase |
| **17** | SDG84795.1 | 351 | Histidinol-phosphate transaminase |
| **18** | SDG84811.1 | 68 | MbtH family protein |
| **19** | SDG84830.1 | 311 | Winged helix-turn-helix transcriptional regulator |
| **20** | SDG84854.1 | 283 | transposase |

**Table S5: PRISM and antiSMASH predictions for NRPS (genes 15 and 16).** Score is not available for antiSMASH predictions (NA=not available).

| | ***PRISM Predictions*** | | | | ***antiSMASH Predictions*** |
|---|---|---|---|---|---|
| **A1** | Val | Ile | Phe | Ser | Val |
| Score | 547.9 | 464.3 | 453.6 | 437.6 | NA |
| **A2** | dPhe | dLeu | dGly | dAla | dLeu |
| Score | 486.6 | 486.5 | 456.7 | 455.2 | NA |
| **A3** | Phe | Trp | β-methyl-Phe | Lys | ?-Phe |
| Score | 563.6 | 492.3 | 479.1 | 474.6 | NA |
| **A4** | dOrn | dEnd | dArg | dLys | ?-dOrn |
| Score | 447.4 | 444.5 | 440.3 | 406.3 | NA |
| **A5** | Thr | Ser | 2,3-dehydroaminobutyric acid | Phe | Thr |
| Score | 791 | 562.5 | 501.8 | 495.8 | NA |
| **A6** | dEnd | dArg | dPhe | dOrn | ?-dPhe |
| | 513.9 | 437.5 | 389.1 | 380.5 | NA |
